# A molecular switch modulates assembly and host factor binding of the HIV-1 capsid

**DOI:** 10.1101/2022.08.25.505312

**Authors:** Randall T. Schirra, Nayara F. B. dos Santos, Kaneil K. Zadrozny, Iga Kucharska, Barbie K. Ganser-Pornillos, Owen Pornillos

**Author notes:** Equal contribution (Randall T. Schirra, Nayara F. B. dos Santos). The Peter Gilgan Centre for Research and Learning, Hospital for Sick Children, Toronto, Ontario, Canada (Iga Kucharska).

## Abstract

Upon entry into a new host cell, the HIV-1 capsid performs multiple essential functions, which include shielding the genome from innate immune sensors^1^, promoting reverse transcription^2^ and transporting the core from the entry site at the plasma membrane to the integration site inside the nucleus^3,4^. The HIV-1 capsid is a fullerene cone made of hexamers and pentamers of the viral CA protein^5,6^. The two types of capsomers are quasi-equivalent, with the same structural elements mediating distinct inter-subunit contacts. In other studied quasi-equivalent viruses, the capacity of genetically identical subunits to form hexamers and pentamers is conferred by molecular switches. Such a switch has not been previously found in retroviral CA proteins. Here, we report cryoEM structures of the HIV-1 CA pentamer within assembled in vitro capsids at nominal resolutions of 2.4-3.4 Å. Comparison with the hexamer identified an internal loop that adopts distinct conformations, 3_10_ helix in the pentamer and random coil in the hexamer. Designed manipulations of the coil/helix configuration allowed us to control pentamer and hexamer formation in a predictable manner, thus proving its function as a molecular switch. Importantly, the switch controls not only fullerene cone assembly, but also the capsid’s capacity to bind post-entry host factors that are critical for viral replication. Furthermore, the switch forms part of the binding site of the new ultra-potent HIV-1 inhibitor, lenacapavir. These studies reveal that a critical assembly element also controls the post-assembly functions of the capsid, and provide new insights on capsid inhibition and uncoating.

## Main

The fullerene cone architecture of the HIV-1 capsid represents an extreme case of quasi-equivalence, in which essentially each of the 1,200 or more copies of the viral CA protein occupies a different chemical environment and thus adopts a different structural configuration^5–7^. Along the body of the cone, the CA subunits are assembled on a hexagonal lattice, with the CA N-terminal domain (NTD) making hexameric rings that are linked together by the C-terminal domain (CTD). In this part of the capsid, conformational variability afforded by CA’s two-domain organization accounts for quasi-equivalence, which manifests in the continuously changing curvature of hexagonal lattice along the body of the cone^6,8^. Twelve CA pentamers occupy sharp points of curvature – termed declinations – exactly 12 of which are required for the capsid to completely close^5^. The molecular rules that dictate CA hexamer or pentamer formation have not been previously eludicated. Furthermore, the relative contributions of CA hexamers and pentamers to HIV-1 capsid interactions with host factors in the post-entry pathway have also been unknown. To address these key gaps, we performed cryoEM and biochemical analyses of the HIV-1 CA pentamer and its complexes with host factors that modulate capsid stability, promote nuclear import, and direct integration site targeting.

### Preparation and single particle averaging analysis of conical HIV-1 capsids in vitro

The classic in vitro model system for the HIV-1 capsid are tubular capsid-like particles (CLPs), which assemble when purified HIV-1 CA protein is incubated in high salt concentrations (≥1 M NaCl) and basic pH^7,9^. These tubes are made only of CA hexamers. Recently, the cellular metabolite inositol hexakisphosphate (IP6) was shown to stabilize the HIV-1 capsid, by binding to the central channel of the CA hexamer^10,11^. IP6 also induces assembly of CA in vitro, even at physiological salt conditions^10^. We found that the IP6-induced CLPs have a range of morphologies – capped tubes and cones – when assembled at pH 8, but are primarily conical at pH 6 (Extended Data Fig. 1). These observations indicate that IP6 promotes incorporation of CA pentamers into the assembling lattice, allowing formation of declinations that close the capsid shell. By adapting deep 2D classification particle selection^12^ and lattice-mediated alignment strategies^13^, we determined the structure of the declination from projection images of the IP6-induced conical CLPs (Fig. 1a,b, Extended Data Fig. 2a, Extended Data Table 1). Local refinement with imposed five-fold symmetry resulted in a high-quality map at a nominal resolution of 3.4 Å, in which the central pentamer and surrounding hexamers are well-defined (Fig. 1c, Extended Data Fig. 2d,e, Movie 1). Both the pentamer (Fig. 1d) and hexamer (Fig. 1e) are similar to their corresponding lower-resolution “in situ” structures that were determined by sub-tomogram averaging of capsids in intact virions^6,14^ (Extended Data Fig. 3). Our results establish that the IP6-induced HIV-1 CA assemblies are the closest in vitro structural mimic of the HIV-1 capsid now available.

**Figure 1.**
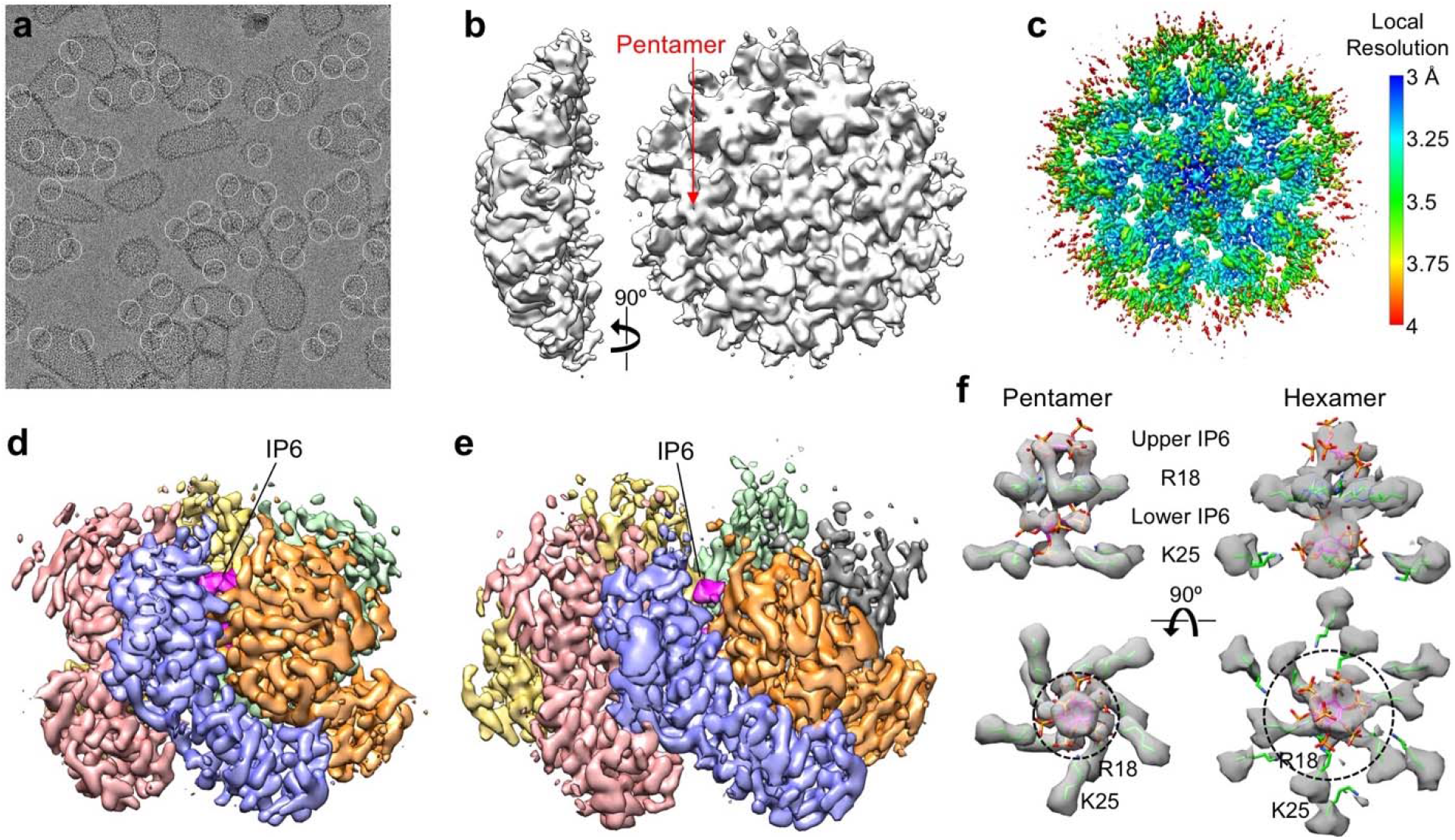
Assembly and structural analysis of IP6-induced HIV-1 CA capsid-like particles (CLPs). **a**, In vitro assembled CLPs, with declinations encircled. **b**, Initial ab initio map calculated from 670,480 particles. **c**, Map after further local refinement with imposed five-fold symmetry, colored according to local resolution. At this contour level (0.4), the central pentamer and the three closed hexamer subunits are very well defined. **d**, Structure of the pentamer (0.28 contour). Each subunit is in a different color, and IP6 is colored in magenta. **e**, Structure of the hexamer (0.22 contour). **f**, Side views (upper) and bottom views (lower) of the central channels of the capsomers. Maps (gray) are contoured at 0.4 (pentamer) and 0.21 (hexamer). Green sticks indicate the Arg18 and Lys25 sidechains. IP6 molecules were rigid-body docked as the *myo* isoform. Dashed circles with indicated diameters circumscribe the positions of the Lys25 epsilon amines.

### IP6 stabilizes the pentameric HIV-1 CA ring

Both the hexamer and pentamer contain two IP6 molecules inside their central channels: one above and one below the ring of positively-charged Arg18 sidechains; the lower IP6 molecules are also coordinated by a second ring of Lys25 sidechains (Fig. 1f). These support the model that charge neutralization promotes formation of the hexamer and pentamer. However, removal of the charges have different effects on CA assembly in vitro. Whereas the R18A substitution abolishes IP6-dependent assembly of CA altogether^10^, the K25A mutant remains competent in assembling tubes^15^. These observations suggest that Arg18 interactions stabilize both the hexamer and pentamer, whereas Lys25 interactions are more important for the pentamer. Consistent with this interpretation, the pentamer channel is narrower at the position of the Lys25 ring (dashed circles in Fig. 1f), allowing close, direct contacts between the primary amines of the lysine sidechains and the phosphates of the lower IP6 molecule.

### Quasi-equivalence of the HIV-1 CA hexamer and pentamer

In quasi-equivalent assembly systems, formation of different oligomers by the same protein is controlled by a molecular switch, which adopts different configurations that are evident from comparison of high-resolution structures of the quasi-equivalent states^16^. Superposition of the HIV-1 CA subunits in the hexamer and pentamer (Fig. 2a-b), allowed us to identify a putative switch in the NTD, comprising a conserved Thr58-Val59-Gly60-Gly61 (TVGG) motif that adopts two alternative configurations – random coil in the hexamer and 3_10_ helix in the pentamer (Fig. 2c). These configurations are well-defined in our cryoEM map (Extended Data Fig. 2e, Movie 2). The interpretation that the TVGG motif functions as a switch is supported by its structural context: the motif mediates alternative packing of the NTD-NTD and NTD-CTD interfaces that hold together the capsomers (Fig. 2d,g,j).

**Figure 2.**
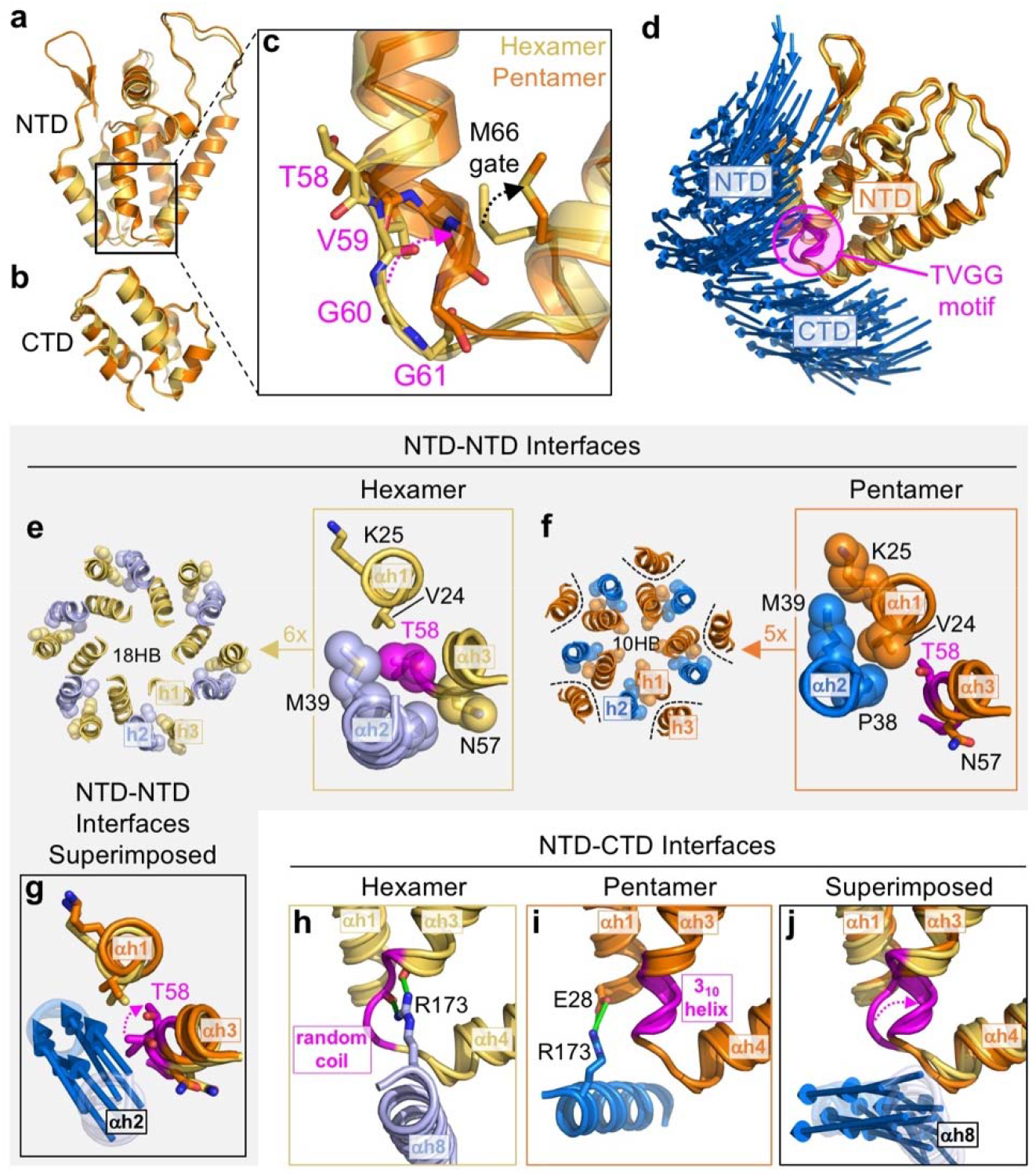
Comparison of HIV-1 CA pentamer and hexamer structures identify a molecular switch. (**a-b**) The NTD and CTD from hexameric (yellow) and pentameric (orange) subunits are superimposed. (**c**) The Thr58-Val59-Gly60-Gly61 (TVGG) motif that connects helices 3 and 4 in the NTD adopt distinct secondary structures in the hexamer and pentamer. (**d**) The TVGG motif (magenta) is located at the juncture of the NTD-NTD and NTD-CTD interfaces that hold together both capsomers. One NTD (pentamer in orange and hexamer in yellow) are superimposed for reference. Two groups of displacement vectors (blue arrows) indicate significant changes in the positions of the NTD and CTD of the neighboring subunit. (**e-g**) Details of the NTD-NTD interfaces in the hexamer (yellow and light blue) and pentamer (orange and blue). Dashed magenta arrow in **g** signifies overwinding of Thr58. Displacement vectors (blue arrows) signify “slippage” of helix 2 from the hexamer state into the pentamer. (**h-j**) Details of the NTD-CTD interfaces in the hexamer (yellow and light blue) and pentamer (orange and blue). Green indicates hydrogen bonds. Dashed magenta arrow in **j** signifies refolding of the TVGG motif. Displacement vectors (blue arrows) indicate movement of helix 8.

In the hexamer, the NTD-NTD interface consists of a 3-helix repeating unit (helix 2 of one subunit and helices 1 and 3 of the adjacent subunit) that makes an 18-helix barrel (Fig. 2e). In the pentamer, helix 3 is excluded from the repeating unit, which then makes a 10-helix barrel (Fig. 2f). Refolding of the TVGG motif facilitates switching from the 3-helix to the 2-helix unit through a ratcheting mechanism. In the hexamer, Thr58 together with Asn57 are the C-terminal residues of helix 3 and engage in reciprocal knobs-in-holes packing against Pro38 and Met39 in helix 2 of the neighboring subunit (Fig. 2e, boxed). In the pentamer, the protein backbone overwinds at Thr58 to facilitate the α-to-3_10_ helix transition (Fig. 2f, boxed). Although the conformational change of this residue is subtle, it seems sufficient to disfavor the knobs-in-holes packing of helices 2 and 3, and instead Pro38-Met39 now packs against Val24-Lys25 in helix 1. Thus, analogous to the “pawl” of a ratchet, Pro38-Met39 in helix 2 engages two alternative “gears”: Asn57-Thr58 in helix 3 to form the hexameric NTD-NTD interface or Val24-Lys25 in helix 1 to form the pentameric interface. Refolding of the TVGG motif toggles alternative packing by remodeling the hexamer gear, allowing the pawl to “slip” into the pentamer state (Fig. 2g).

The TVGG motif also mediates alternative packing of the NTD-CTD interface (Fig. 2h-j). In the hexamer, helix 8 of the CTD is adjacent to the helix 3/4 loop (Fig. 2h). The highly conserved Arg173 residue in this helix makes two hydrogen bonds: one that caps the C-terminal end of helix 3 (Asn57) and another to the carbonyl of Val59 (green in Fig. 2h). These are part of an extensive hydrogen bonding network (Extended Data Fig. 4a) that we previously showed to facilitate pivoting of the CTD to accommodate the variable curvature of the hexagonal capsid lattice^17^. Our results suggest an additional function in stabilizing both the random coil configuration of the switch and the α-helical character of helix 3. In the pentamer, the helix 3/4 loop with the 3_10_ helix configuration no longer interacts with helix 8 due to repositioning of the CTD (Fig. 2i). Instead, Arg173 makes hydrogen bonds with a different partner (Glu29) in helix 1 (green in Fig. 2i), and the C-terminal end of helix 7 (NTD) and portions of the NTD-CTD linker pack against the groove between helices 8 and 11 (CTD) (Extended Data Fig. 4a).

In summary, the TVGG motif within the helix 3/4 loop of HIV-1 CA adopts two distinct secondary structural folds – random coil and 3_10_ helix – each of which engages a distinct set of quaternary packing interactions in the hexamer and pentamer. The location of the motif allows it to simultaneously modulate both the NTD-NTD and NTD-CTD interfaces (Fig. 2d,g,j), explaining how these two sets of interactions cooperate during capsid assembly.

Additional, more subtle conformational changes further distinguish the pentameric state of NTD subunit from its hexameric state. Refolding of the TVGG loop is accompanied by a change in the rotamer configuration of Met66, which we describe here to function as a gate (Fig. 2c). In the hexamer state, the Met66 gate is in a “closed” configuration and must move into an “open” configuration in order to prevent clash with the folded 3_10_ helix in the pentamer state. Refolding of the TVGG switch also remodels the hydrophobic core of the NTD. In the hexamer, the NTD core contains a large hidden pocket surrounded by hydrophobic sidechains (Extended Data Fig. 4b). This pocket is smaller in the pentamer, because the empty space is partly occupied by the repositioned Val59 sidechain. Finally, the Ala31-Phe32 peptide bond in the helix 1/2 loop rotates by 180° compared to the hexamer (Extended Data Fig. 4c). The Phe32 sidechain points towards the hydrophobic core of the protein and lines the hidden pocket in both states. These additional elements likely form part of the allosteric network that triggers TVGG switching during assembly.

Importantly, all of the residues identified above are important for viral replication or inhibition. Mutations in the TVGG motif generally result in non-viable or severely impaired viruses^18–21^. Mutations in the Pro38-Met39 “pawl” in helix 2 abolishes CA assembly or highly destabilizes the capsid^22,23^. Asn57 and the Met66 gate modulate binding of the assembled capsid to host factors important for nuclear import and integration, as well as binding to one class of capsid-targeting inhibitors^24–33^. The hidden pocket in the NTD core that is walled by Phe32 and Val59 is also a binding site for a different class of inhibitors^18,34^. Altogether, the TVGG motif is positioned at a critical site for capsid function and inhibition, indicating that it may have post-assembly functions as well.

### Designed manipulations of the TVGG switch predict HIV-1 CA assembly

To test our hypothesis that the TVGG motif is a molecular switch, we designed structure-based mutations to control the coil/helix configuration, which should predict the in vitro assembly behavior of the CA protein and resulting CLP morphology (Fig. 3a). Solution conditions can also affect CLP morphology (Extended Data Fig. 1), and so for uniformity we assayed for assembly using the IP6-induced conditions at pH 6, which in our hands produces conical CLPs with wild type (WT) CA (Fig. 3b, Extended Data Fig. 1c).

**Figure 3.**
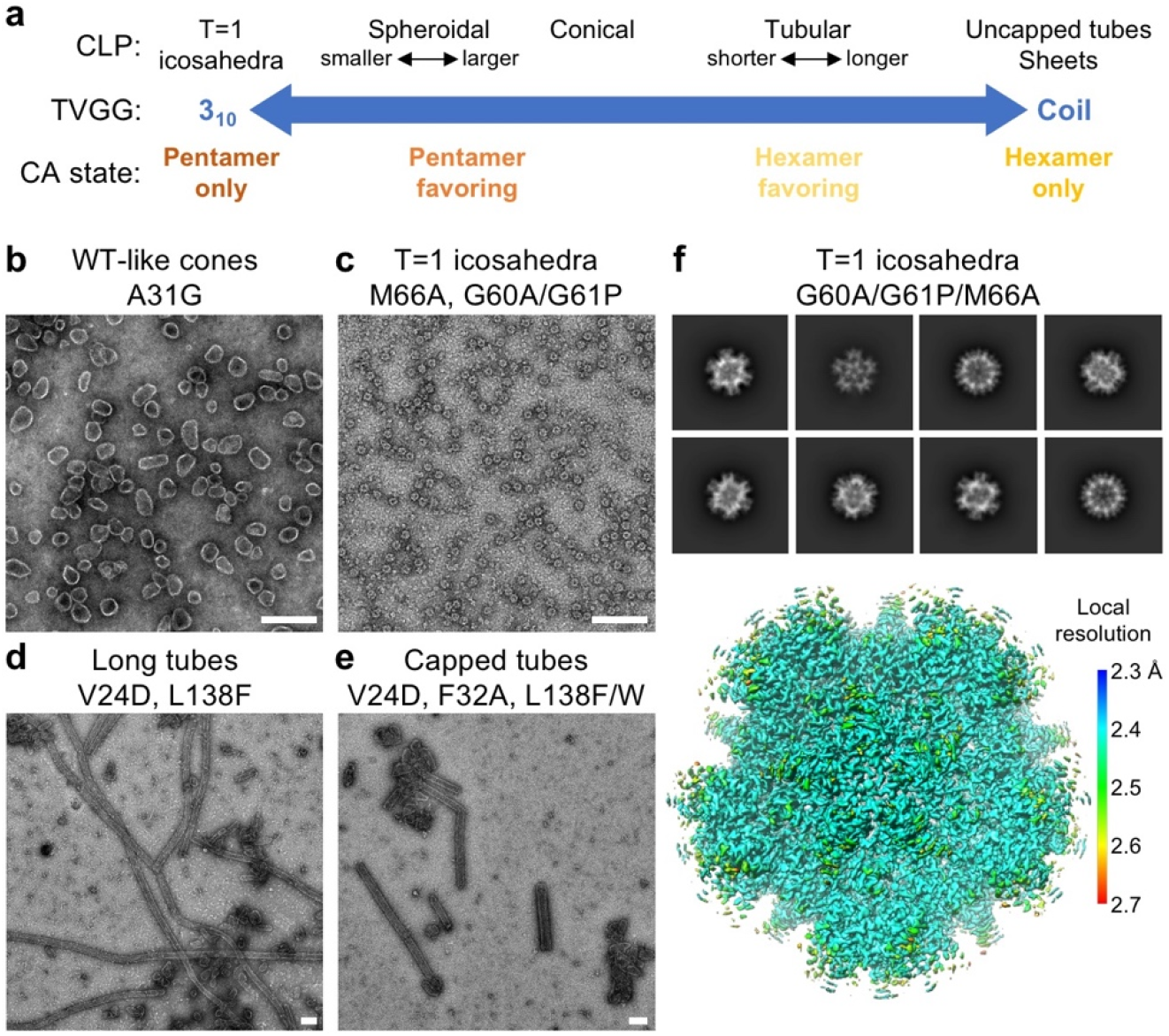
Designed manipulations favor or disfavor pentamer formation of HIV-1 CA. (**a**) Predicted phenotypes of CA assembly behavior in vitro and CLP shape, as determined by the configuration of the TVGG motif. All but T=1 icosahedra have been observed previously. (**b-e**) Morphologies of CLPs formed by the indicated CA mutants in 25 mM MES, pH 6, 150 mM NaCl, 5 mM β-mercaptoethanol, 4 mM IP6. Scale bars, 200 nm. (**f**) 2D class averages (upper) and cryoEM structure (lower) of T=1 icosahedra made by CA G60A/G61P/M66A. The map is colored according to local resolution.

WT HIV-1 CA is known to favor hexamers in vitro, implying that the coil-to-helix transition of the TVGG motif is rate limiting. We surmised that the Met66 gate must constitute part of the activation barrier for switching from hexamer to pentamer. Thus, a CA mutant with a constitutively open gate (M66A) should favor pentamer switching. Indeed, M66A only formed small spheres of around 20 nm in diameter (Fig. 3c, Extended Data Fig. 1d). CryoEM imaging indicated that the spheres are T=1 particles, or icosahedra made of 12 pentamers and no hexamers. Next, we replaced the two glycines in the TVGG motif with alanine and proline, since our structural analysis indicated that these two residues would rigidify and stabilize the 3_10_ helix configuration. The G60A/G61P mutant also assembled into T=1 particles, as did the G60A/G61P/M66A triple mutant (Fig. 3c, Extended Data Fig, 1e,f). We solved the structure of the triple mutant to 2.4 Å resolution (Fig. 3f, Extended Data Fig. 2c-d, Extended Data Table 1), which confirmed that the pentamer has the same structure as the WT pentamer, and that the switch is in the 3_10_ helix configuration (Extended Data Fig. 2f). We next asked if this mutant can still form hexamers by assembling it in 1 M NaCl at pH 8, which induces only tubes with WT CA (Extended Data Fig. 1a). The triple mutant only formed aggregates in these conditions (Extended Data Fig. 1f). Our interpretation is that the G60A/G61P/M66A mutations locked the CA protein in its pentamer state.

We were unable to identify amino acid substitutions to rigidify the random coil configuration of the TVGG switch, which we predicted would have locked CA in its hexamer state. Nevertheless, structure-based mutations can be introduced to disfavor switching into the pentamer. As described above, Val24 stabilizes the pentameric NTD ring by mediating the knobs-in-holes interaction of helices 1 and 2 (Fig. 3d). Accordingly, the V24D mutation favored assembly of long, tubular CLPs when incubated with IP6 at pH 6 (Fig. 3d, Extended Data Fig. 1g). We also surmised that modifying the loosely-packed hydrophobic core of the NTD might inhibit repacking of the internal pocket and thus pentamer switching. To increase the packing density without modifying the TVGG motif, we replaced Leu138 with larger sidechains; this residue is in helix 7, across the internal pocket from Val59 (Extended Data Fig. 4b). The L138F, L138W and L138Y mutations significantly destabilized CA. Nevertheless, L138F and L138W still formed cones, but had higher proportions of tubular CLPs compared to WT (Fig. 3d-e, Extended Data Fig. 1h-j). Finally, we tested the importance of the conformational change in the helix 1/2 loop involving Ala31 and Phe32 (Extended Data Fig. 4c). As with the Leu138 substitutions, A31G and F32A also destabilized the protein and reduced assembly efficiency, yet still supported cone formation (Extended Data Fig. 1j-k). F32A had a higher proportion of tubular capsids, again indicating that pentamer formation is disfavored (Extended Data Fig. 1k). All of these pentamer-disfavoring (or hexamer-favoring) mutations we tested did not completely abolish CA’s ability to make pentamers, likely because they do not rigidify the TVGG switch in its random coil configuration.

### The HIV-1 CA hexamer/pentamer switch modulates ligand binding

One of the critical functions of the HIV-1 capsid is to recruit host cell factors that promote viral replication in terminally-differentiated T cells and macrophages, the natural hosts of HIV-1. Because the details of these interactions are typically studied biochemically and structurally in context of the HIV-1 CA hexamer, and because genetic approaches cannot distinguish between the hexamer and pentamer, any potential contributions from the pentamer have been unknown. A major class of such capsid-binding factors include CPSF6, NUP153 and SEC24C, which contain one or more phenylalanine-glycine (FG) motifs that bind within the NTD-CTD interface of the hexamer and make contacts with both the NTD and CTD^26,27,29,33^. Binding of these natural FG ligands is competitively inhibited by pharmacological inhibitors, PF74^26–28^ and GS-CA1/GS-6207/lenacapavir^30–32^, each of which contains an analogous phenyl ring. The FG-motif phenylalanine ring binds to a hydrophobic pocket in the NTD that is demarcated by the helix 3/4 loop containing the TVGG switch. This NTD pocket alone is sufficient to allow weak binding of FG motif-containing peptides and inhibitors^24,25^. Tighter binding is afforded in context of the hexamer by additional contacts with the CTD, which adopts a “closed” configuration and contacts ligand moieties outside of the phenyl ring^26,27^. While it was suggested that the alternative configurations of the CTD relative to the NTD might modulate ligand binding^6^, our structures indicate a direct mechanism of inhibition, because refolding of the TVGG motif also remodels the phenylalanine pocket (Fig. 4a). Specifically, the open configuration of the Met66 gate in the pentamer would generate a steric clash with the phenyl ring, which is critical for binding. We experimentally tested this prediction by incubating the IP6-induced WT CLPs with excess amounts of an FG-motif peptide derived from CPSF6. This peptide was previously shown to bind the NTD alone^25^ as well as the hexameric NTD-CTD interface^26,27^. We solved the structure of the declination in these capsid-peptide complexes to a nominal resolution of 5.0 Å (Extended Data Fig. 2b,d, Extended Data Table 1). Because the structure contains both the pentamer and its surrounding hexamers, the hexamers serve as an internal positive control, and indeed bound peptides were clearly seen as U-shaped densities within all of the hexameric NTD-CTD interfaces (Fig. 4b). We did not find peptide densities within the pentameric NTD-CTD interfaces (Fig. 4c).

**Figure 4.**
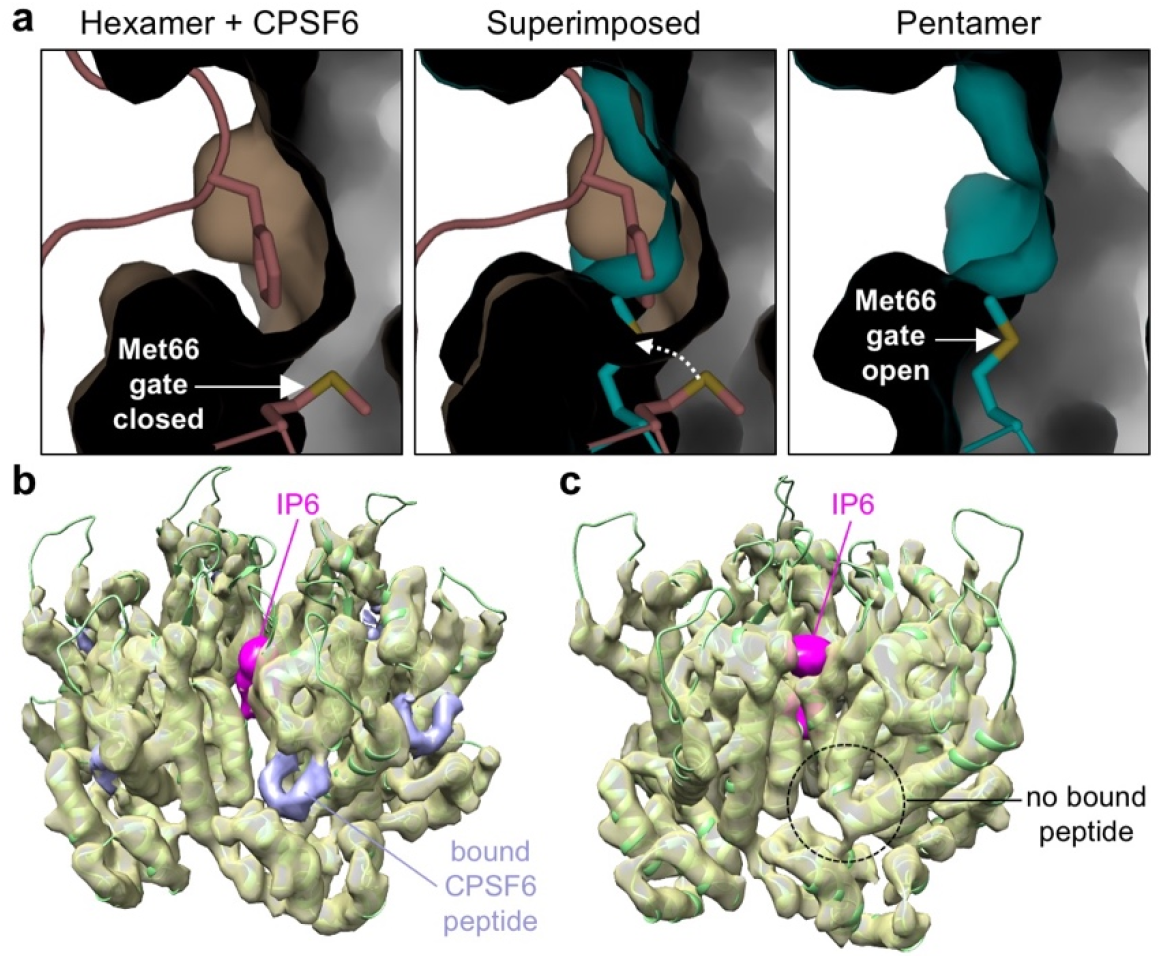
Conformational switching of the TVGG motif remodels the FG-motif binding site. (**a**) Cutaway views of the phenylalanine pocket in a CPSF6-bound X-ray crystal structure of the CA hexamer (PDB 4wym; left) and our cryoEM structure of the pentamer (right). The CA protein is shown in surface representation. Configurations of the Met66 gate are indicated. Dashed white arrow in the middle panel signifies gate opening (hexamer to pentamer). (**b-c**) Views of the hexamer and pentamer from the cryoEM reconstruction (5.0 Å resolution) of the declination from conical CLPs incubated with excess CPSF6 FG peptide. Pink indicates bound IP6, found in both hexamer and pentamer. Lavender indicates that CPSF6 is only bound to the hexamer.

## Discussion

Molecular switches are triggered by allostery. We find here that the HIV-1 CA switch is an internal loop at the juncture of two major assembly interfaces. This central location allows the switch to directly and simultaneously modulate the CA NTD-NTD and NTD-CTD interactions, and reveals how allosteric communication and cooperativity between these separate sets of contacts are generated during capsid assembly. Interestingly, our results also suggest that the allosteric network that triggers switching includes the protein hydrophobic core, because altering residues that line the core (Phe32 and Leu138, as well as the Met66 gate) modulates switching into the pentamer. This phenomenon has been documented in protein kinases, in which hydrophobic core residues move synchronously to act as a non-covalent bridge between the regulatory and catalytic “splines” of the enzyme located on opposite sides of the protein fold^35^. We propose that CA’s helix 1/2 (NTD-NTD interface) and helix 3/4 (NTD-NTD and NTD-CTD interfaces) are analogous “splines” bridged by the residues that line its loosely packed hydrophobic core. Allosteric inhibition of kinases has been successfully exploited pharmacologically^36,37^, and similar strategies may also be applicable to HIV-1 CA.

Another important insight from these studies is our discovery that the HIV-1 CA hexamer/pentamer switch controls not just the assembly process, but also the capacity of the assembled capsid to bind FG-motif host cell factors that are important for the post-entry pathway of the virus. Our results indicate that the nuclear factor CPSF6, and likely also NUP153 and SEC24C, bind only to the hexamer. This finding reinforces the importance of the assembled capsid lattice in the nuclear import process. Other studies indicate that pentamer-specific factors might facilitate core trafficking or docking to the nucleus^3,38^, pointing to division of labor between the two capsomers and supporting the requirement for an intact or nearly intact capsid at these viral replication steps.

While it may be that the switch configuration irreversibly locks the binding capacity of the capsomers in the assembled capsid, a more intriguing possibility is that ligand binding might trigger its conformational transition, post-assembly. Back-switching of the TVGG motif from 3_10_ helix to its coil configuration could be the basis of the proposed capsid remodeling during passage of the viral core through the nuclear pore channel^39,40^ (which is densely packed with FG-motifs), or alternatively, the basis of capsid disassembly or uncoating. Importantly, the new ultra-potent HIV-1 inhibitor, lenacapavir, binds to the same site on the capsid as FG-motifs. Indeed, a single point mutation in the methionine gate (M66I) confers resistance to this drug, although at severe cost to viral fitness^31^; the basis of this fitness cost can now be explained by our findings. We speculate that induced-fit binding of lenacapavir to pentamer subunits may trigger back-switching, which could destabilize the capsid and induce premature uncoating. In support of this idea, even though lenacapavir and similar compounds stabilize tubes made of CA hexamers^24,27,30^, they promote fracturing of the fullerene cone capsid^2,41,42^.

## Methods

### Protein purification and assembly

HIV-1 CA proteins were purified as described^43^. Conical CLPs were assembled by incubating the protein (12 mg/mL) in 4 mM IP6, 50 mM MES, pH 6, 150 mM NaCl, 5 mM β-mercaptoethanol at 37 °C for 1-2 h. Tubular CLPs were assembled by incubating the protein in 50 mM Tris, pH 8, 1 M NaCl, 5 mM β-mercaptoethanol at 37 °C for 1-2 h. CA mutants were purified and assembled in the same way as the WT protein. Peptide-bound WT CLPs were prepared by incubation (30 min on ice) of assembled CA (in IP6) with 3-fold molar excess of synthetic CPSF6-FG peptide (GTPVLFPGQPFGQPPLG, N-terminally acetylated and C-terminally amidated; Celltein).

### CryoEM sample preparation and data collection

Conical CLPs were diluted 8 to 12-fold with 25 mM MES, pH 6, 100 μM IP6 and then immediately applied (4 μL) to glow-discharged lacey carbon 300-mesh Cu grids. T=1 particles were purified away from unassembled protein by using size exclusion chromatography (Superdex 200 10/300 GL; Cytiva) in 25 mM MES, pH 6, 50 mM NaCl, 10 mM β-mercaptoethanol, 100 μM IP6, and 4 μL of relevant fractions were applied to C-flat 1.2/1.3 300-mesh holey carbon grids. WT CLPs bound to CPSF6 were prepared in the same way as the unbound CLPs. Grids were briefly blotted manually, and plunge-frozen in liquid ethane using a home-built device.

CryoEM data were collected at the University of Virginia Molecular Electron Microscopy Core (MEMC), using a Titan Krios (ThermoFisher) operating at 300 kV and equipped with an energy filter and K3 direct detector (Gatan). For the WT CLPs and T=1 particles, videos were collected in EPU (ThermoFisher) at a pixel size of 1.08 Å in counting mode, with a total dose of 50 electrons/Å^2^ over 40 frames and target defocus of −0.5 to −2.5 μm. For CPSF6-bound CLPs, videos were collected at a pixel size of 1.62 Å, with a total dose of 54 electrons/Å^2^.

All image processing and map calculations were performed in cryoSPARC v.3.3.1^44^ (Extended Data Table 1). Raw movies were corrected for beam-induced motion using MotionCor2^45^ and CTF estimation was performed with CTFFIND4^46^, as implemented in cryoSPARC. Initial particles were manually picked to generate references for subsequent template-based picking.

For the WT CA declination structure, 13,439,772 particles were initially extracted. Two rounds of reference-free 2D classification and selection were performed to identify and remove junk particles (mostly carbon edges). The selected 3,985,778 particles were then processed through several rounds of iterative reference-free 2D classification and selection, resulting in 1,345,054 particles that were used for *ab initio* structure calculation (5 models, C1 symmetry). Four models (1,075,923 particles) converged on similar maps and were combined into a single set through one round of refinement with C1 symmetry and low-pass filter of 20 Å. The resulting map was then rotated and centered on the pentamer. Heterogeneous refinement (3D classification, 2 models) was then performed in C1, with a low-pass filter of 20 Å. After discarding overlaps, the final set of 525,219 particles were refined with C5 symmetry and local CTF estimation, resulting in a map at a nominal resolution of 3.4 Å.

For the G60A/G61P/M66A T=1 structure, 694,288 particles were extracted. After two rounds of reference-free 2D classification, 509,666 particles were used for *ab initio* calculation with I symmetry imposed. Homogeneous refinement resulted in a final map at a nominal resolution of 2.4 Å.

For the CPSF6-bound structure, 5,655,057 initial particles were processed for removal of junk particles, resulting in 1,168,289 particles that were iteratively classified and selected using reference-free 2D alignment. After one round of heterogeneous refinement (using as reference the WT structure filtered to 20 Å), the final set of 198,306 particles were refined with C5 symmetry to a nominal resolution of 5.0 Å.

Coordinate models were built for the WT declination and T=1 particle, by docking PDB 4xfx into the maps, followed by iterative manual re-building in Coot^48^ and refinement using Phenix^47^ (phenix.real_space_refine). For the declination, refinements were performed separately for two groups: the pentamer subunit (chain A) and the three closest three hexamer subunits to the pentamer (chains L, M and N) (Extended Data Table 1). Non-crystallographic symmetry restraints were used when appropriate. The remaining three hexamer subunits (chains O, P and Q) were modeled by rigid-body docking of the final refined model of chain M as polyalanine. For the T=1 particle, an initial round of morphing was performed prior to real space refinement. In both maps, IP6 densities were 5-fold symmetric and so modeling was restricted to rigid-body fitting of the *myo* isoform as described for the hexamer^10^.

### Negative stain EM

Samples (4 μL) were applied to Formvar continuous carbon 300-mesh Cu grids and incubated for 2 min. Grids were moved to a 20-μL drop of 0.1 M KCl, incubated for 2 min, and blotted. Grids were then placed on a 20-μL drop of 2% uranyl acetate, incubated for 2 mins, and blotted to dryness. Images were collected on a Tecnai Spirit (ThermoFisher) or an F20 microscope (ThermoFisher), both operating at 120 kV.

### Deposition

CryoEM maps are deposited at the Electron Microscopy Data Bank under accession numbers EMD-26715 (WT declination), EMD-26718 (T=1 G60A/G61P/M66A) and EMD-26811 (WT declination bound to CPSF6 peptide). Coordinates are deposited at the Protein Data Bank under accession numbers 7urn (WT declination) and 7urt (T=1).

## Acknowledgements

Transmission electron micrographs were recorded at the University of Virginia Molecular Electron Microscopy Core facility, which was built in part with NIH grant G20-RR31199. Purchase of the Titan Krios was funded in part by S10-RR025067. This study was funded by R21-AI167756 and P50-AI150464 (O.P. and B.K.G.-P.). R.T.S. was supported by T32-GM008136. We especially thank K. Dryden and M. Purdy for invaluable assistance in EM data collection and processing, and M. Purdy for help with optimizing server and software set up.

## Contributions

Cloning, mutagenesis, protein purification and assembly: R.T.S., N.F.B.d.S., K.K.Z, I.K. Negative stain EM: R.T.S., N.F.B.d.S., B.K.G.-P. CryoEM sample preparation and data collection: R.T.S., N.F.B.d.S., B.K.G.-P. Image data processing: R.T.S., B.K.G.-P., O.P. Coordinate modeling and refinement: R.T.S., O.P. Project conceptualization and funding acquisition: B.K.G.-P. and O.P. Protocol design and development: R.T.S., N.F.B.d.S., I.K., B.K.G.-P., O.P. Manuscript preparation: O.P. with input from all authors.

---

**Extended Data Table 1.**
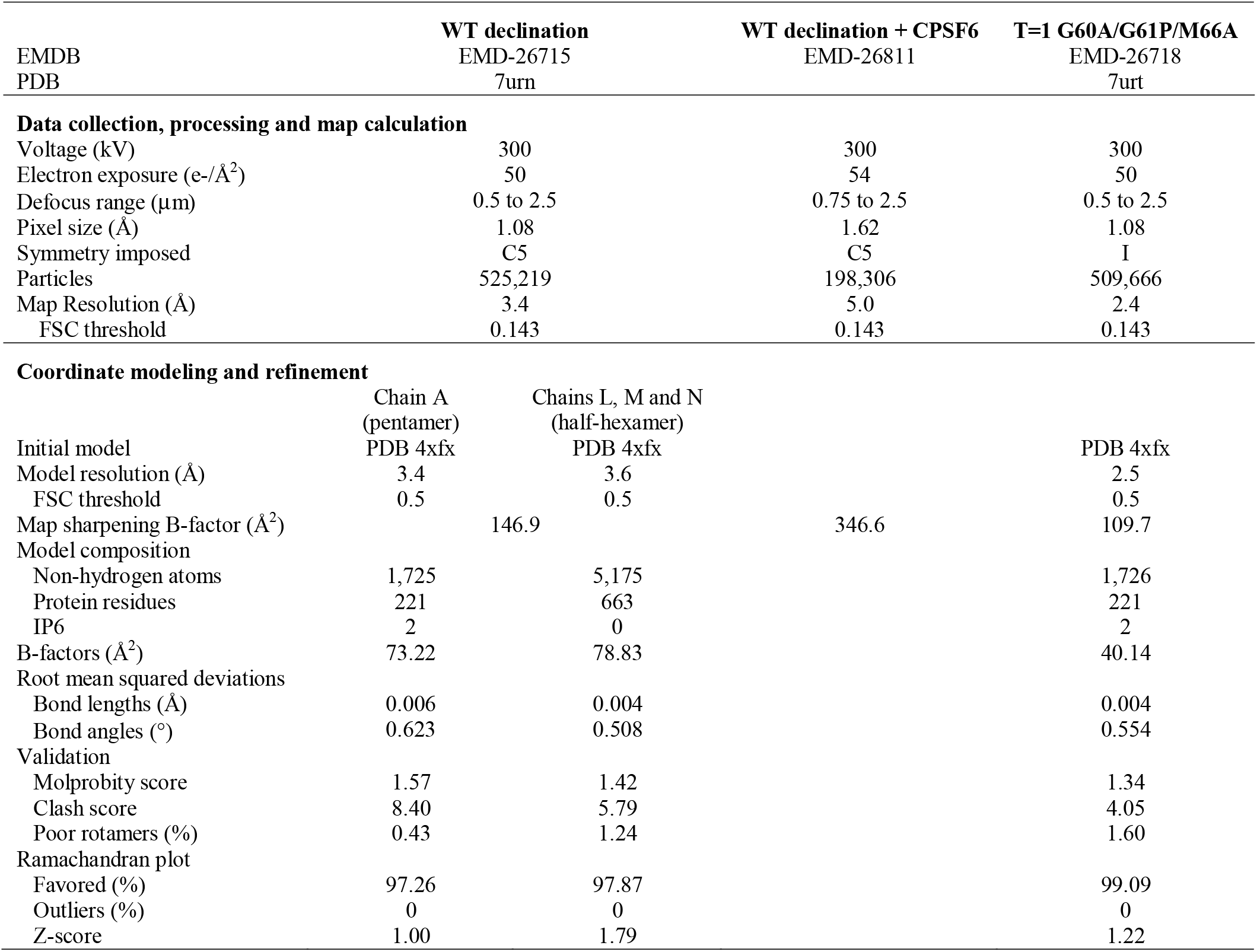
Data collection, processing and refinement statistics

**Extended Data Figure 1.**
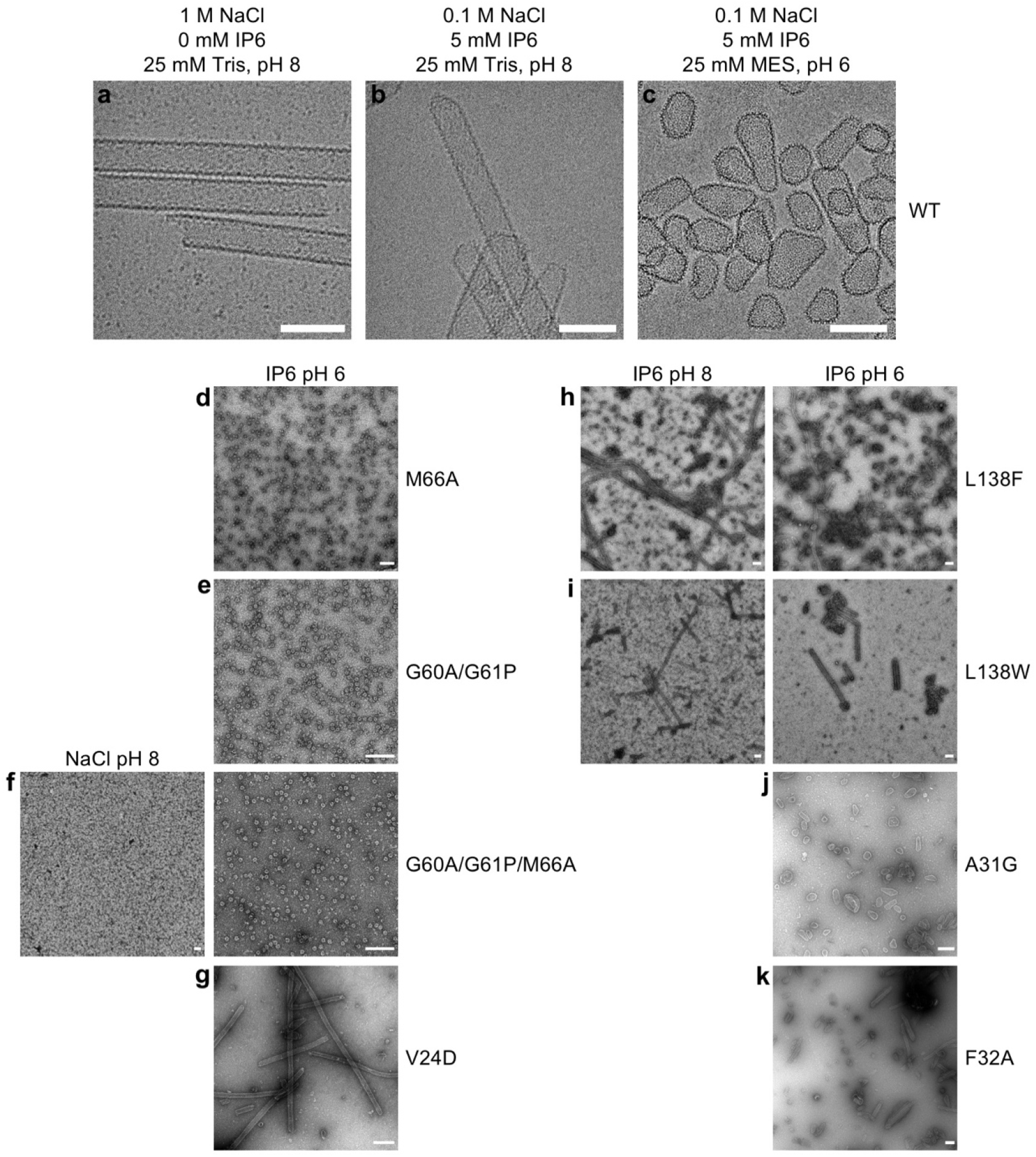
**a-c**, Cryoimages of WT HIV-1 CA CLPs assembled under the indicated conditions. Reactions were incubated at 37 °C for 1 h. Scale bars, 100 nm. **d-k**, Negative stain images of CLPs formed by the indicated CA mutants, under the indicated conditions. Scale bars, 200 nm.

**Extended Data Figure 2.**
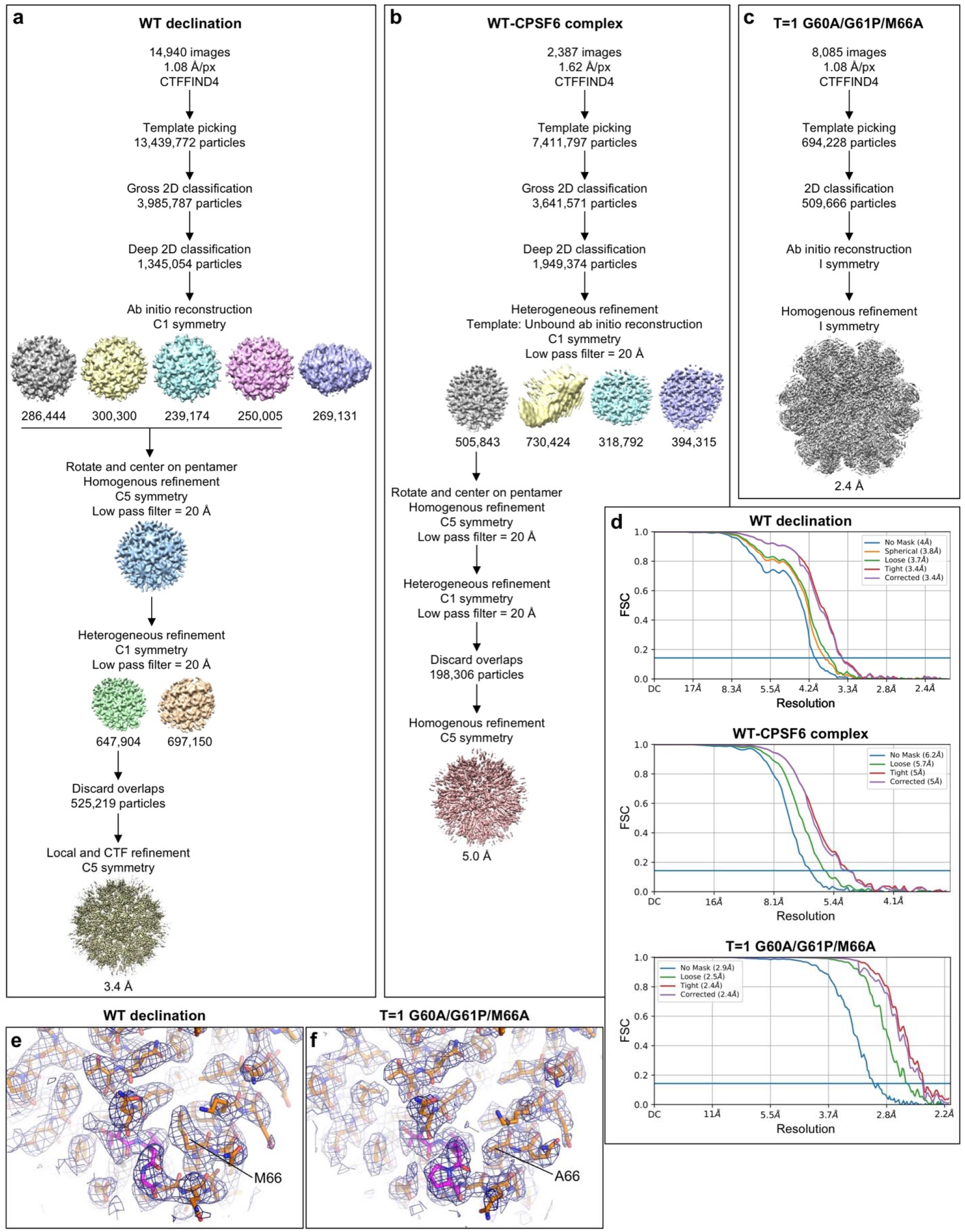
**a-c**, Single particle averaging workflows for the indicated structures. **d**, Fourier shell correlation (FSC) plots from final refinement rounds. **e**, Map (blue mesh) and real-space-refined model (sticks) of the WT pentamer. The TVGG motif is in magenta. **f**, Map (blue mesh) and real-space-refined model (sticks) of the G60A/G61P/M66A pentamer. The mutated TVAP motif is in magenta.

**Extended Data Figure 3.**
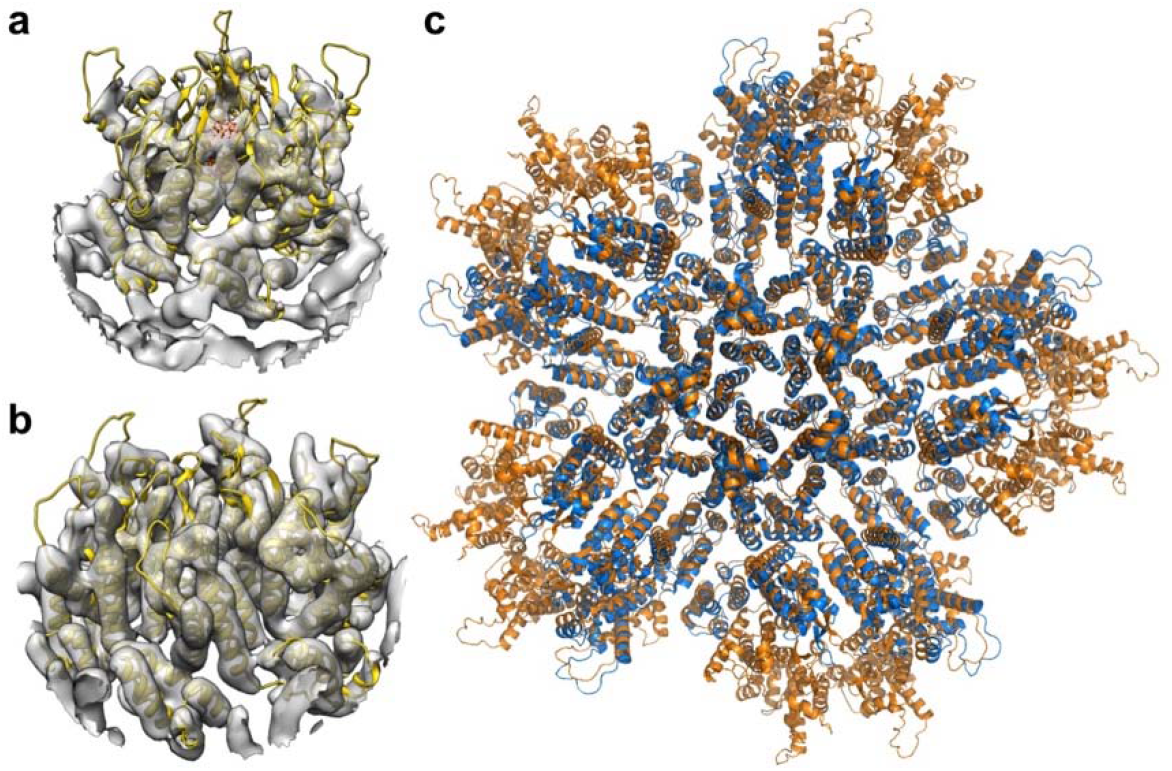
Structures of the capsomers in IP6-induced in vitro CLPs are similar to their corresponding structures in situ. **a**, Docking of our real-space-refined CA pentamer model (yellow ribbons, with IP6 in sticks) into EMD 3466 (gray), the first reported in situ structure of the HIV-1 CA pentamer at 8.8 Å resolution. **b**, Docking of our real-space-refined CA hexamer model (yellow ribbons) into EMD 3465 (gray), the in situ structure of the HIV-1 CA hexamer at 6.8 Å resolution. **c**, Superposition of the final PDB model of the declination after real space refinement into our 3.4 Å map (orange) with PDB 5mcy (blue).

**Extended Data Figure 4.**
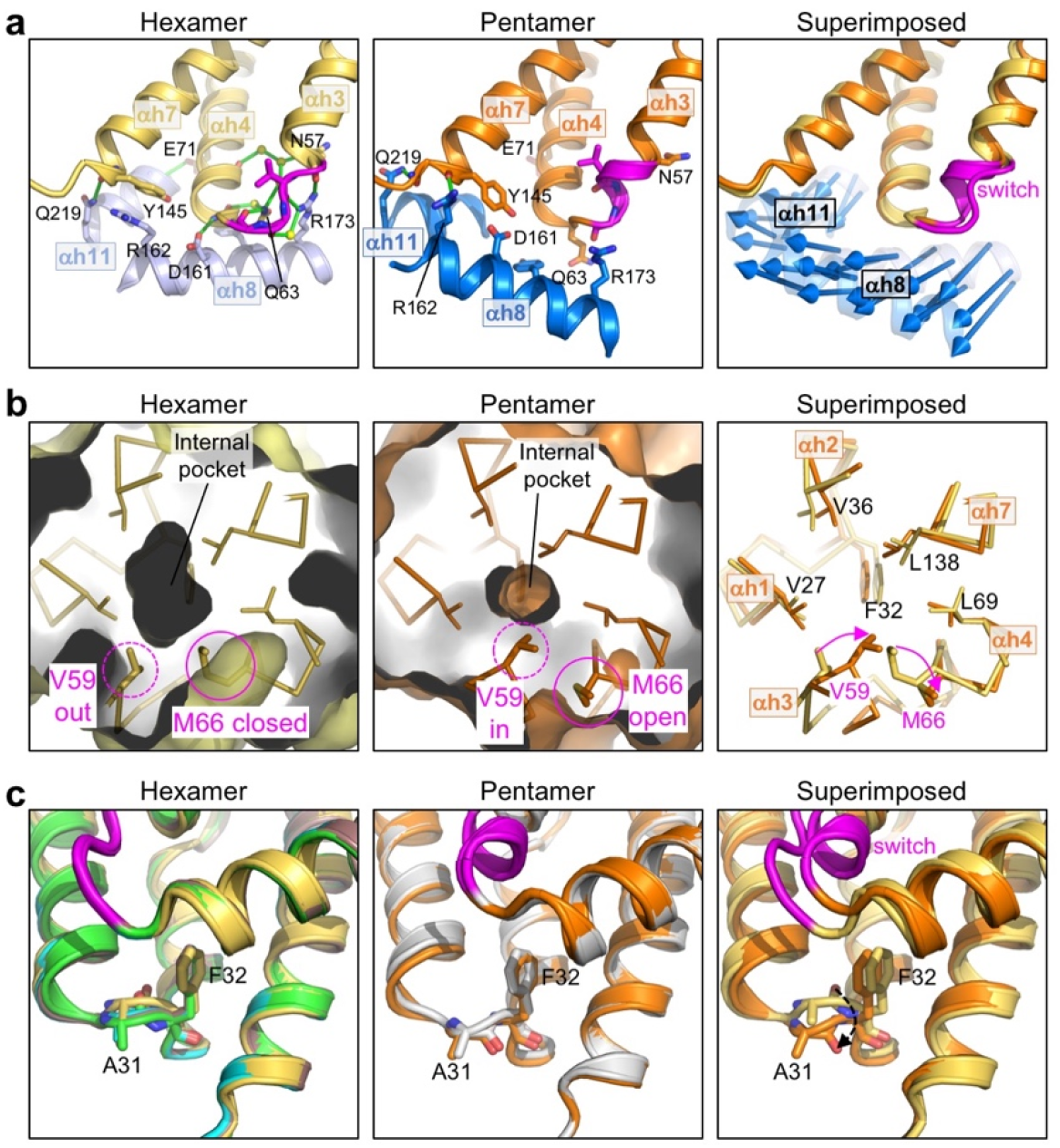
Additional distinguishing structural features of the HIV-1 CA hexamer and pentamer. **a**, The NTD-CTD interfaces are distinguished by different sets of hydrogen bonds. Left: The hexameric interface is stabilized by an extensive network of hydrogen bonds involving the TVGG motif, surrounding residues and ordered water molecules. Shown is PDB 3h47, a high-resolution (1.9 Å) X-ray crystal structure. Middle: In the pentamer, inter-subunit hydrogen bonds involving the TVGG motif are absent. Arg162 and Gln219 make hydrogen bonds with the C-terminal end of helix 7 and the linker, respectively. Right: Blue arrows indicate the displacement vectors for each residue in helices 8 and 11. **b**, The NTD hydrophobic core is underpacked. The hexamer pocket (Left) is larger than the pentamer pocket (Middle). Right: Residues lining the pocket are shown as sticks and labeled. **c**, Rotation of the Ala31-Phe32 peptide bond. Left: Hexamer structures, with PDB 3h47 in yellow. The three best-defined hexamer subunits in our WT CA declination structure are shown (chain L, green; chain M, cyan; chain N, brown). Middle: Pentamer structures of WT CA (orange, 3.4 Å) and G60A/G61P/M66A (gray, 2.4 Å). Right: PDB 3h47 is superimposed with the WT pentamer. Dashed arrow indicates rotation of the peptide bond.

## References

1 Hilditch, L. & Towers, G. J. A model for cofactor use during HIV-1 reverse transcription and nuclear entry. Curr Opin Virol 4, 32–36, doi:10.1016/j.coviro.2013.11.003 (2014).

2 Christensen, D. E., Ganser-Pornillos, B. K., Johnson, J. S., Pornillos, O. & Sundquist, W. I. Reconstitution and visualization of HIV-1 capsid-dependent replication and integration in vitro. Science 370, abc8420, doi:10.1126/science.abc8420 (2020).

3 Zila, V., Muller, T. G., Muller, B. & Krausslich, H. G. HIV-1 capsid is the key orchestrator of early viral replication. PLoS Pathog 17, e1010109, doi:10.1371/journal.ppat.1010109 (2021).

4 Naghavi, M. H. HIV-1 capsid exploitation of the host microtubule cytoskeleton during early infection. Retrovirology 18, 19, doi:10.1186/s12977-021-00563-3 (2021).

5 Ganser, B. K., Li, S., Klishko, V. Y., Finch, J. T. & Sundquist, W. I. Assembly and analysis of conical models for the HIV-1 core. Science 283, 80–83, doi:10.1126/science.283.5398.80 (1999).

6 Mattei, S., Glass, B., Hagen, W. J., Kräusslich, H. G. & Briggs, J. A. The structure and flexibility of conical HIV-1 capsids determined within intact virions. Science 354, 1434–1437, doi:10.1126/science.aah4972 (2016).

7 Li, S., Hill, C. P., Sundquist, W. I. & Finch, J. T. Image reconstructions of helical assemblies of the HIV-1 CA protein. Nature 407, 409–413, doi:10.1038/35030177 (2000).

8 Pornillos, O., Ganser-Pornillos, B. K. & Yeager, M. Atomic-level modelling of the HIV capsid. Nature 469, 424–427, doi:10.1038/nature09640 (2011).

9 Ehrlich, L. S., Agresta, B. E. & Carter, C. A. Assembly of recombinant human immunodeficiency virus type 1 capsid protein in vitro. J Virol 66, 4874–4883, doi:10.1128/JVI.66.8.4874-4883.1992 (1992).

10 Dick, R. A. et al. Inositol phosphates are assembly co-factors for HIV-1. Nature 560, 509–512, doi:10.1038/s41586-018-0396-4 (2018).

11 Mallery, D. L. et al. IP6 is an HIV pocket factor that prevents capsid collapse and promotes DNA synthesis. Elife 7, e35335, doi:10.7554/eLife.35335 (2018).

12 Yao, X., Fan, X. & Yan, N. Cryo-EM analysis of a membrane protein embedded in the liposome. Proc Natl Acad Sci U S A 117, 18497–18503, doi:10.1073/pnas.2009385117 (2020).

13 Briggs, J. A. et al. Structure and assembly of immature HIV. Proc Natl Acad Sci U S A 106, 11090–11095, doi:10.1073/pnas.0903535106 (2009).

14 Ni, T. et al. Structure of native HIV-1 cores and their interactions with IP6 and CypA. Sci Adv 7, eabj5715, doi:10.1126/sciadv.abj5715 (2021).

15 Renner, N. et al. A lysine ring in HIV capsid pores coordinates IP6 to drive mature capsid assembly. PLoS Pathog 17, e1009164, doi:10.1371/journal.ppat.1009164 (2021).

16 Domitrovic, T. et al. Virus assembly and maturation: auto-regulation through allosteric molecular switches. J Mol Biol 425, 1488–1496, doi:10.1016/j.jmb.2013.02.021 (2013).

17 Pornillos, O. et al. X-ray structures of the hexameric building block of the HIV capsid. Cell 137, 1282–1292, doi:10.1016/j.cell.2009.04.063 (2009).

18 Lemke, C. T. et al. Distinct effects of two HIV-1 capsid assembly inhibitor families that bind the same site within the N-terminal domain of the viral CA protein. J Virol 86, 6643–6655, doi:10.1128/JVI.00493-12 (2012).

19 Manocheewa, S., Swain, J. V., Lanxon-Cookson, E., Rolland, M. & Mullins, J. I. Fitness costs of mutations at the HIV-1 capsid hexamerization interface. PLoS One 8, e66065, doi:10.1371/journal.pone.0066065 (2013).

20 Rihn, S. J. et al. Extreme genetic fragility of the HIV-1 capsid. PLoS Pathog 9, e1003461, doi:10.1371/journal.ppat.1003461 (2013).

21 Al-Mawsawi, L. Q. et al. High-throughput profiling of point mutations across the HIV-1 genome. Retrovirology 11, 124, doi:10.1186/s12977-014-0124-6 (2014).

22 Forshey, B. M., von Schwedler, U., Sundquist, W. I. & Aiken, C. Formation of a human immunodeficiency virus type 1 core of optimal stability is crucial for viral replication. J Virol 76, 5667–5677, doi:10.1128/jvi.76.11.5667-5677.2002 (2002).

23 Ganser-Pornillos, B. K., von Schwedler, U. K., Stray, K. M., Aiken, C. & Sundquist, W. I. Assembly properties of the human immunodeficiency virus type 1 CA protein. J Virol 78, 2545–2552, doi:10.1128/jvi.78.5.2545-2552.2004 (2004).

24 Blair, W. S. et al. HIV capsid is a tractable target for small molecule therapeutic intervention. PLoS Pathog 6, e1001220, doi:10.1371/journal.ppat.1001220 (2010).

25 Price, A. J. et al. CPSF6 defines a conserved capsid interface that modulates HIV-1 replication. PLoS Pathog 8, e1002896, doi:10.1371/journal.ppat.1002896 (2012).

26 Price, A. J. et al. Host cofactors and pharmacologic ligands share an essential interface in HIV-1 capsid that is lost upon disassembly. PLoS Pathog 10, e1004459, doi:10.1371/journal.ppat.1004459 (2014).

27 Bhattacharya, A. et al. Structural basis of HIV-1 capsid recognition by PF74 and CPSF6. Proc Natl Acad Sci USA 111, 18625–18630, doi:10.1073/pnas.1419945112 (2014).

28 Gres, A. T. et al. X-ray crystal structures of native HIV-1 capsid protein reveal conformational variability. Science 349, 99–103, doi:10.1126/science.aaa5936 (2015).

29 Buffone, C. et al. Nup153 Unlocks the Nuclear Pore Complex for HIV-1 Nuclear Translocation in Nondividing Cells. J Virol 92, doi:10.1128/JVI.00648-18 (2018).

30 Yant, S. R. et al. A highly potent long-acting small-molecule HIV-1 capsid inhibitor with efficacy in a humanized mouse model. Nat Med 25, 1377–1384, doi:10.1038/s41591-019-0560-x (2019).

31 Link, J. O. et al. Clinical targeting of HIV capsid protein with a long-acting small molecule. Nature 584, 614–618, doi:10.1038/s41586-020-2443-1 (2020).

32 Bester, S. M. et al. Structural and mechanistic bases for a potent HIV-1 capsid inhibitor. Science 370, 360–364, doi:10.1126/science.abb4808 (2020).

33 Rebensburg, S. V. et al. Sec24C is an HIV-1 host dependency factor crucial for virus replication. Nat Microbiol 6, 435–444, doi:10.1038/s41564-021-00868-1 (2021).

34 Kelly, B. N. et al. Structure of the antiviral assembly inhibitor CAP-1 complex with the HIV-1 CA protein. J Mol Biol 373, 355–366, doi:10.1016/j.jmb.2007.07.070 (2007).

35 Kim, J. et al. A dynamic hydrophobic core orchestrates allostery in protein kinases. Sci Adv 3, e1600663, doi:10.1126/sciadv.1600663 (2017).

36 Zhang, J., Yang, P. L. & Gray, N. S. Targeting cancer with small molecule kinase inhibitors. Nat Rev Cancer 9, 28–39, doi:10.1038/nrc2559 (2009).

37 Dar, A. C. & Shokat, K. M. The evolution of protein kinase inhibitors from antagonists to agonists of cellular signaling. Annu Rev Biochem 80, 769–795, doi:10.1146/annurev-biochem-090308-173656 (2011).

38 Santos da Silva, E. et al. HIV-1 capsids mimic a microtubule regulator to coordinate early stages of infection. EMBO J 39, e104870, doi:10.15252/embj.2020104870 (2020).

39 Blanco-Rodriguez, G. et al. Remodeling of the Core Leads HIV-1 Preintegration Complex into the Nucleus of Human Lymphocytes. J Virol 94, doi:10.1128/JVI.00135-20 (2020).

40 Guedan, A. et al. HIV-1 requires capsid remodelling at the nuclear pore for nuclear entry and integration. PLoS Pathog 17, e1009484, doi:10.1371/journal.ppat.1009484 (2021).

41 Shi, J., Zhou, J., Shah, V. B., Aiken, C. & Whitby, K. Small-molecule inhibition of human immunodeficiency virus type 1 infection by virus capsid destabilization. J Virol 85, 542–549, doi:10.1128/JVI.01406-10 (2011).

42 Marquez, C. L. et al. Kinetics of HIV-1 capsid uncoating revealed by single-molecule analysis. Elife 7, e34772, doi:10.7554/eLife.34772 (2018).

43 Yoo, S. et al. Molecular recognition in the HIV-1 capsid/cyclophilin A complex. J Mol Biol 269, 780–795, doi:10.1006/jmbi.1997.1051 (1997).

44 Punjani, A., Rubinstein, J. L., Fleet, D. J. & Brubaker, M. A. cryoSPARC: algorithms for rapid unsupervised cryo-EM structure determination. Nat Methods 14, 290–296, doi:10.1038/nmeth.4169 (2017).

45 Zheng, S. Q. et al. MotionCor2: anisotropic correction of beam-induced motion for improved cryo-electron microscopy. Nat Methods 14, 331–332, doi:10.1038/nmeth.4193 (2017).

46 Rohou, A. & Grigorieff, N. CTFFIND4: Fast and accurate defocus estimation from electron micrographs. J Struct Biol 192, 216–221, doi:10.1016/j.jsb.2015.08.008 (2015).

47 Liebschner, D. et al. Macromolecular structure determination using X-rays, neutrons and electrons: recent developments in Phenix. Acta Crystallogr D Struct Biol 75, 861–877, doi:10.1107/S2059798319011471 (2019).

48 Emsley, P., Lohkamp, B., Scott, W. G. & Cowtan, K. Features and development of Coot. Acta Crystallogr D Biol Crystallogr 66, 486–501, doi:10.1107/S0907444910007493 (2010).

